# The impact of the soluble protein fraction and kernel hardness on wheat flour starch digestibility

**DOI:** 10.1101/2022.02.14.480385

**Authors:** Jia Wu, Frederick J. Warren

## Abstract

Wheat is the staple crop for 35% of the world’s population, providing a major source of calories for much of the world’s population. Starch is the main source of energy in wheat flour, but the digestibility of wheat starch varies greatly between different flours and wheat products. This has relevance from a health perspective because wheat starch products that are rapidly digested and elicit large post-prandial glucose peaks are associated with a host of cardiac and metabolic disorders. In this study, we investigate the impact of protein on starch digestion in three commercially sourced flours with different grain hardness. Grain hardness impacted on flour particle size, but not significantly on starch digestion. A soluble extract of wheat proteins was found to dramatically reduce starch digestion, even following gastric proteolysis. Proteomic analysis revealed that this soluble extract was enriched in proteinaceous α-amylase inhibitors which were partially degraded during gastric proteolysis. Therefore, we conclude that the soluble proteins of wheat flour have a significant contribution towards retarding starch digestion, even following gastric digestion.

## 1. Introduction

Starch is one of the most abundant sources of polysaccharides widely distributed in seeds, roots and tubers of plants, which supports about half of the carbohydrate caloric intake for much of the population of the planet (Jobling, 2004; Whistler & Daniel, 2000). Amongst starchy staple crops, one of the most widely consumed is bread wheat (*Triticum aestivum*), and understanding the factors which impact wheat starch digestion has the potential to yield nutritional benefit (Shewry, Hazard, Lovegrove, & Uauy, 2020). Starchy foods vary greatly in their digestibility in the gastrointestinal tract, and starchy foods which are rapidly digested can result in large peaks in post-prandial blood glucose concentration (Brouns et al., 2008). Regular consumption of foods which produce large excursions in post-prandial glycaemia, including wheat based products such as white bread, are has been associated with health problems such as insulin resistance, hyperinsulinemia, obesity, cardiovascular diseases and type 2 Diabetes (Borneo & León, 2012; Lanzerstorfer et al., 2018). Thus, reducing the digestion rate of starch in wheat based foods is a potential route to reduce post-prandial glycaemic responses (Brouns et al., 2008; Eelderink, Schepers, Preston, Vonk, Oudhuis, & Priebe, 2012).

Foods made from bread wheat are generally made from the milled endosperm of the wheat grain (although whole wheat and other products may include other parts of the grain). The resulting flour is mainly starch (80-85%), but also contains other components which can impact on the digestibility of the starch, including protein, which comprises 10-15% of bread wheat flour. The main proteins in bread wheat are the gluten proteins, glutenin and gliadin, which are storage proteins that comprise ~80% of the protein in bread flour (Juhász, Békés, & Wrigley, 2015). These proteins form the largely insoluble gluten network which give the typical properties of many wheat based foods such as bread. Bhattarai, Dhital, and Gidley (2016) reported an increase of starch digestibility after the removal of gluten in wheat flour, indicating that gluten may play a role in modulating starch digestibility through acting as a barrier to digestive enzymes. Zou et al. (2019) further confirmed that the presence of a gluten network (in this case in durum wheat based products) could cause a reduction of α-amylase activity even after cooking with boiling water. Zou et al. (2019) further highlighted 12 different thermally stable proteinaceous α-amylase inhibitors present in wheat which they speculated may also contribute to the overall decrease of enzyme activity.

Low molecular weight (~12KDa) proteinaceous α-amylase inhibitors ((α-AIs) are the next most abundant protein in the starchy endosperm after gluten proteins, and may be present in monomeric, dimeric or trimeric forms. Plant α-AIs have been reported to be capable of reducing digestibility of starch, lowering blood sugar level and enhancing insulin responses in *in vivo* and *in vitro* studies (Alagesan, Raghupathi, & Sankarnarayanan, 2012; López-Barón, Gu, Vasanthan, & Hoover, 2017; Sales, Souza, Simeoni, Magalhães, & Silveira, 2012; Svensson, Fukuda, Nielsen, & Bønsager, 2004). Kodama et al. (2005) showed reductions of postprandial glycaemic levels in subjects who were served wheat albumin (the soluble fraction of wheat proteins including α-AI’s) in addition to normal meals. Other proteins, including puroindolines in the albumin fraction of wheat proteins may also impact on starch digestibility indirectly through impacting on starch damage during milling (Juhász et al., 2015). However, the relative contribution of these different protein families to starch digestion in wheat based foods is not fully understood. A further challenge is that to impact significantly on starch digestion in the small intestine, wheat proteins must escape proteolysis in the stomach. L. Li, Fan, and Zhao (2022) investigated proteinaceous α-AIs extracted from different plants by analysing their inhibition activities following treatment with different temperature, pH and digestive proteinases. Inhibition activities of the extracts were decreased under certain conditions, especially after treating with pepsin. Therefore, the effectiveness of different protein components to reduce starch digestion will be further impacted by gastric digestion.

In this study we aimed to elucidate the impact of protein fractions within wheat flour on the digestibility of starch. Using commercially available flours with different grain hardness and protein contents we determined the impact of gastric protein digestibility on starch digestibility using an *in vitro* model system. Proteomic analysis was used to identify the soluble proteins present in the digestion mixture before and after gastric digestion. We demonstrate that, while grain hardness and insoluble proteins had a relatively minor impact on starch digestibility, the soluble albumin fraction containing high levels of proteinaceous α-AI had a significant impact on starch digestibility, even following extensive gastric digestion. The role of wheat proteinaceous α-AI should be considered a significant factor impacting wheat starch digestibility in future studies.

## 2. Materials and Methodology

### 2.1 Materials

Baker’s Green wheat flour (BKG, protein content: 12.5% w/w), Titan wheat flour (Titan, protein content: 12.4% w/w, high in gluten) and Heygates plain all-purpose wheat flours (HG, protein content: 10.0% w/w) were kindly supplied as a gift by Mervyn Poole (Heygates Limited Northampton, UK). The Titan and BKG flours were both hard wheats with a grain hardness of 46-80 single-kernel characterization system (SKCS) units, while the HG flour was a soft flour with a grain hardness of 15-46 SKCS units (information on grain hardness kindly supplied by Mervin Poole, Heygates Limited). The control wheat starch was obtained from Tereos Syral (Aalst, Belgium). Samples were all received as milled powders, and no additional milling steps were applied in this study. Porcine stomach pepsin (EC 3.4.23.1) was purchased from ThermoFisher, Waltham, US with a stock activity of 800-2500 U/mg protein. Pancreatic α-amylase (A6255) was purchased from Sigma/Merck, Darmstadt, Germany with a stock activity of 1095 U/mg protein.

### 2.2 Starch extraction from wheat flours

Pre-milled wheat flours were suspended in water at a ratio of 1 to 12, then the pH of slurry was adjusted to 8.5 with 0.1 M NaOH. The slurry was mixed for 5 minutes and then centrifuged at 3000 rpm for 10 minutes. The viscous top layer was removed, and the rest of the precipitate was re-suspended into deionised water. After mixing, the resuspended slurry was centrifuged again to remove top layers. The washing process was repeated two more times, then all the supernatant was removed. The resulting white precipitate was then suspended in deionised water and sequentially filtered through a 300 μm sieve and a 53 μm sieve. Starch extracted from wheat flours was collected after air drying.

### 2.3 Starch content analysis for wheat flours

Total starch contents (TTS) of each of the wheat flours was analysed using the total starch assay kit (K-TSTA-100A, Megazyme, Ireland) (B. McCleary, T. Gibson, & D. Mugford, 1997; B. V. Mccleary, T. S. Gibson, & D. C. Mugford, 1997). The rapid total starch (RTS) method was applied in this study, as described by the manufacturer. Briefly, 100 mg of each wheat flour was firstly incubated with thermostable α-amylase in boiling water bath for 15 minutes, then further degraded with amyloglucosidase in a 50 °C water bath for 30 minutes. Those hydrolysates were analysed using the GOPOD method to determine the glucose concentration and this was then converted to the total starch value (%, w/w). Triplicates of each sample were analysed.

### 2.4 Starch digestion by using pancreatic α-amylase following pepsin treatment

#### 2.4.1 Endpoint starch digestion

This experiment consisted of two digestive steps: Firstly, samples were incubated with pepsin (using a modification of Englyst et al. (2018)). Control wheat starch, three wheat flours, and extracted wheat starch were weighed to give a sample equivalent to 100 mg of starch and individually transferred into 15 ml conical tubes. Deionised water (1 ml) and pepsin solution (2 ml, 5 mg/ml in 0.05 M HCL) were added into each sample tube. A control group (no pepsin) was prepared similarly by adding 1 ml of deionised water and 2 ml of 0.05 M HCL to individual pre-weighed samples. Then all the tubes were incubated in a 37 °C water bath for 30 minutes. After incubation, the pH of the digestion solution was neutralised by adding 2 ml of 0.05 M NaOH and the volume was made up to 10 ml with 5 ml of phosphate buffer (PBS). 2. The resulting solutions were further digested with pancreatic α-amylase (method was modified from Zou, Sissons, Gidley, Gilbert, and Warren (2015))All sample tubes were then placed on a rotator (60 rpm) in a 37 °C incubator for 5 minutes to calibrate temperature. Time zero controls were prepared before additions of α-amylase by taking 100 μl of digestion solution from each sample tube and transferring into microcentrifuge tubes (1.2 ml capacity) containing 100 μl of 0.3 M Na_2_CO_3_ (stop solution). Starch digestion was started by adding 100 μl of α-amylase solution (20 U/ml in reaction mixture) into each sample tube. Then, the tubes were incubated at 37 °C with end-over-end mixing at 60 rpm for 2 hours. Starch digestion was stopped at 2 hours by transferring 100 μl digestion solution from sample tube into microcentrifuge tubes tubes containing 100 μl of stop solution. Microcentrifuge tubes containing time zero controls and endpoint digestion samples were stored at −18 °C for subsequent determination of maltose release using the PAHBAH method. Each digestion experiment was independently prepared and carried out in triplicate.

#### 2.4.2 Digestion of control wheat starch with addition of wheat flour extracts

Each of the wheat flours was weighted based on 100 mg starch and transferred in to 15 ml conical tubes followed by the addition of 1 ml of deionised water and 2 ml of 0.05 M HCL. The tubes were mixed and then incubated in a 37 °C water bath for 30 minutes. After incubation, 2 ml of 0.05 M NaOH was added into each tube for neutralising pH. PBS (5 ml) was then added to establish 10 ml of solution. The tubes were vortexed for 10 seconds, then centrifuged at 3000 rpm for 10 minutes. The supernatant (9 ml) from each tube was then respectively transferred new tubes containing pre-weighed 100 mg control wheat starch and 1 ml of PBS (the control group was prepared by adding 1 ml of water, 2 ml of 0.05 M HCL, 2 ml of 0.05 M NaOH and 5 ml of PBS into 100 mg control wheat starch). The continue steps for starch digestion with α-amylase were the same as method described in the section 2.4.1. Analysed samples were all prepared in duplicates.

### 2.5 Analysis of reducing sugar released from starch by using PAHBAH method

Reducing sugar in the digesta samples collected in 2.4.1 and 2.4.2 was quantified using the PAHBAH method modified from (Moretti & Thorson, 2008). Microcentrifuge tubes containing digesta samples collected in 2.4.1 and 2.4.2 were defrosted and centrifuged at 3000 rpm for 10 minutes. 10 μl of each supernatant was taken from each microcentrifuge tube and mixed with 190 μl deionised water in a 96 well plate. After mixing, 100 μl of each diluted solution was transferred into a fresh microcentrifuge tube. A dilution series of maltose standards (1000, 500, 250, 125, 60.3 and 0 μM) was prepared in separate microcentrifuge tubes. PAHBAH reagent was prepared by dissolving 250 mg *p*-hydroxybenzoic acid hydrazide (H9882, Sigma/Merck, Darmstadt, Germany) in 4.75 ml 0.5 M HCL, then adding 45 ml 0.5 NaOH. 1 mL of PAHBAH reagent was added to each microcentrifuge tube. The tubes were incubated in a boiling water bath for 5 minutes. Following cooling at room temperature for 5 minutes 200 μl was withdrawn from each tube and transferred to a 96 well plate (CC7672-7596, Starlab, Milton Keynes, UK). The absorbance of each well was read in a plate reader at 405 nm. The maltose reducing equivalent concentration of each sample was calculated using the maltose standard curve and presented as percentage starch digested based on the total starch value of each sample.

### 2.6 Thermal properties of the flours determined using Differential Scanning Calorimetry (DSC)

Wheat flours were accurately weighed (100 mg) into DSC pans (Hastelloy ampoules, TA Instruments Ltd., New Castle, USA) following the addition of 1 ml degassed deionised water. An additional reference pan was prepared containing 1 mL of degassed deionised water. The pans prepared in triplicate were placed into a TA Instruments multicell differential scanning calorimeter (MC-DSC). The instrument was programmed to heat samples from 20 °C to 150 °C at a rate of 1 °C per minute. The slow heating rate was selected to maintain pseudo-steady state conditions during gelatinisation of the starch (Bogracheva, Wang, Wang, & Hedley, 2002). Data was analysed using TA Universal Analysis 2000 software.

### 2.7 Analysis of relative crystallinity for all wheat flours by using X-ray diffraction (XRD)

All three different wheat flours were analysed with the Rigaku Smartlab SE X-ray diffraction system (Cu X-ray tube, Rigaku, Tokyo, Japan) operating at 40 kV – 50 mA. The scattering angles (2*θ, θ* is the Bragg angle) started from 2° to 40° with a changing rate of 0.02°. The acquisition time was 4° per minute. Data was collected with SmartLab Studio II software. Raw data was then further analysed for relative crystallinities by using Origin 2016 software (OriginLab, Northampton, US). The method for relative crystallinity estimation was adapted from Lopez - Rubio, Flanagan, Gilbert, and Gidley (2008). XRD analysis was carried out in duplicate for each sample.

### 2.8 Particle size distribution analysis for wheat flours by using Static Light Scattering (SLS)

Each wheat flour was suspended into deionised water to establish a 10% (w/v) slurry and their particle size distribution was measured with a Beckman Coulter LS 13-320 (Beckman Coulter, Brea, California, US). The pre-set model “Wheat flour in water” was applied for analysis. Each sample was examined in triplicate.

### 2.9 Characterisation of proteinaceous α-amylase inhibitors using Mass Spectrometry and Proteomics

Each wheat flour was weighed into a 15 mL conical tube on a 100 mg starch basis. Samples were then treated with pepsin by adding 1 ml deionised water and 2 ml pepsin solution (described on section 2.4). A pepsin free control group was also prepared as section 2.4 described. After incubation, the pH of the solution was neutralised by adding 2 ml of 0.05 M NaOH. Pepsin treated samples and controls were centrifuged at 3000 rpm for 10 minutes and then 4 ml supernatant was collected as flour extract for proteomic analysis.

Proteins were precipitated from the flour extract with acetone (Nickerson & Doucette, 2020). Then, pellets were resuspended in 2.5% sodium deoxycholate (SDC) with 0.2M EPPS (Merck) buffer pH8, reduced, alkylated and digested with sequencing grade trypsin (Promega) according to standard procedures based on Shevchenko, Tomas, Havli, Olsen, and Mann (2006). The SDC was precipitated by adjusting to 0.5% formic acid, and the peptides were purified from the supernatant using C13 OMIX tips (Agilent). Aliquots were analysed by nanoLC-MS/MS on an Orbitrap Eclipse™ Tribrid™ mass spectrometer coupled to an UltiMate^®^ 3000 RSLCnano LC system (Thermo Fisher Scientific, Hemel Hempstead, UK). The samples were loaded onto a pre-column (Acclaim™ PepMap™ 100 C18, 5 μm, 1×5mm, Thermo) with 0.1% TFA at 15 μl min-1 for 3 min. The trap column was then switched in-line with the analytical column (nanoEase M/Z column, HSS C18 T3, 100 Å, 1.8 μm; Waters, Wilmslow, UK) for separation using the following gradient of solvents A (water, 0.1% formic acid) and B (80% acetonitrile, 0.1% formic acid) at a flow rate of 0.2 μl min-1: 0-3 min 3% B (parallel to trapping); 3-10 min increase B to 7% (curve 4); 10-70 min increase B to 37%; 70-90 min increase B to 55%; followed by a ramp to 99% B and re-equilibration to 3% B. Data were acquired with the following mass spectrometer settings in positive ion mode: MS1/OT: resolution 120K, profile mode, mass range m/z 300-1800, AGC 100%, fill time 50 ms; MS2/IT: data dependent analysis was performed using parallel CID and HCD fragmentation with the following parameters: top20 in IT turbo mode, centroid mode, isolation window 1 Da, charge states 2-5, threshold 1e4, CE = 33, AGC target 1e4, max. inject time 35 ms, dynamic exclusion 1 count, 15 s exclusion, exclusion mass window ±10 ppm.

Peaklists (mgf) were generated from the MS raw files using the Proteowizard tool MSConvert (Chambers et al., 2012). A database search was performed with Mascot Server 2.8 with the following parameters: databases Uniprot_Triticum_20190904.fasta (130677 entries) and the MQcontaminants database (251 sequences), enzyme trypsin, 2 missed cleavages, MS1/MS2 tolerances 6 ppm/0.6 Da, oxidation (M), deamidation (N,Q) and acetylation (protein N-terminus) were set as variable modifications and carbamido-methylation (CAM) of cysteine as fixed modification. The Mascot search results were imported into Scaffold 4.11.0.

Statistical analysis was carried out by using Scaffold 5.0.1. Statistical significance of differences between the control group and gastric group for each wheat flour were assessed by Fisher’s Exact Test (no specific correction method applied) with the significant level of P-value < 0.05.

## 3. Results and discussion

### 3.1 Inclusion of a gastric digestion step increase the digestibility of starch in wheat flour

Starch digestion was carried out for 3 different commercial wheat flours and wheat starch. There were two groups analysed, the gastric group which was treated with 5 mg/ml pepsin solution to mimic the gastric phase, and the control group which was not treated with pepsin. As Figure 1 shows, all 3 types of wheat flours treated with pepsin (Green bars) had significantly higher digestion rates of starch comparing with control group (Blue bars). There was 30.4%, 37.9% and 28.1% starch digested respectively for the BKG, Titan and HG with a gastric digestion stage, but only 8.0%, 10.9% and 5.9% of starch (respectively) were digested for the control group of each flour. Wheat starch showed overall higher digestibility than any of the flour samples, with 44.0% starch digestion for the control and 43.5% for sample with gastric digestion sample. These findings are in agreement with the study of Zou et al. (2015) using durum wheat, who reported increases in starch digestibility for pasta and semolina with pepsin treatment prior to hydrolysis by α-amylase. Pepsin is known as a non-specific proteinase capable of hydrolysing a broad range proteins (Nelson, Lehninger, & Cox, 2008). The differences in digestibility of starch with and without gastric digestion was not observed for the purified wheat starch. Therefore, the results implied that the digestibility of starch in wheat flours was reduced due to the presence of protein.

**Figure 1.**
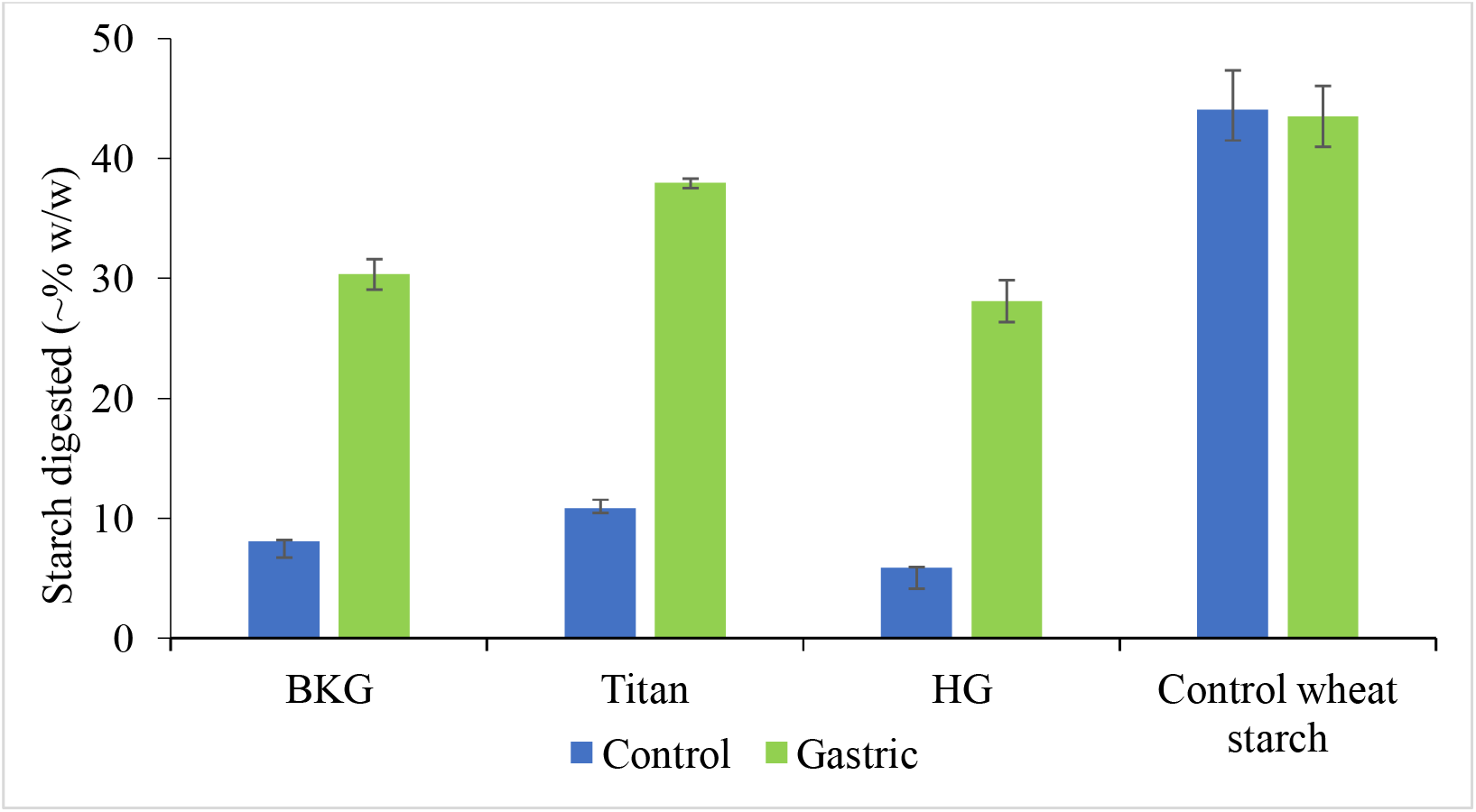
Starch digestion for wheat flours and control wheat starch. Blue bars represent digestions with pancreatic α-amylase alone and the Green bars represent samples treated with pepsin (Gastric phase) prior to pancreatic α-amylase hydrolysis.

Different rates of starch digestion were observed between the different wheat flours for both the gastric and control groups. Titan wheat flour showed the highest digestion rate and the HG showed the lowest, and BKG was in between. Apart from proteinaceous inhibitors for digestive enzymes, there are many other factors to be considered that may affect the digestibility of starch for wheat flours. For example, starch crystallinity, thermal behaviour and particle sizes of flours (Choct & Annison, 2007; Guo, Yu, Copeland, Wang, & Wang, 2018; Heaton, Marcus, Emmett, & Bolton, 1988; Lin et al., 2020; López-Barón et al., 2017; Mahasukhonthachat, Sopade, & Gidley, 2010; Tester, Karkalas, & Qi, 2007). To evaluate the relation of those factors to different digestibility of starch, relative crystallinity, particle size distribution and thermal behaviour was examined for all 3 different wheat flours.

### 3.2 Analysis of particle size distribution, gelatinisation of starch and relative crystallinity for wheat flours

Static light scattering (SLS) was used to analyse particle size distributions for each of the wheat flours. Particle distribution and Mean and Median values are presented in Figure 2. BKG and Titan showed minor differences in size distribution. The particle diameters of Titan (Mean: 92.25 μm, Median 91.25 μm) were slightly smaller than BKG (Mean: 95.84 μm, Median: 95.09 μm). HG was found to have a broader distribution of particle sizes and a larger particle diameter (Mean: 156.6 μm, Median: 153.5 μm). This difference is due to the HG being a soft wheat (15-46 SKCS units), while the BKG and the Titan were hard bread making wheats (46-80 SKCS units), explaining their different milling performances. Differences in particle size might be a potential reason for the previous digestion result which showed the lowest digestion rate of starch for HG, as several studies have indicated that larger particles are more slowly digested (Carré, 2007; de la Hera, Rosell, & Gomez, 2014; Guo, Yu, Wang, Wang, & Copeland, 2018; Lin et al., 2020). Guo, Yu, Wang, et al. (2018) compared starch digestion for raw durum wheat flours milled into different particle size and found that starch digested slower in raw durum wheat flours with larger particle sizes. However, the differences in particle size observed in this study are smaller than in the referenced literature, so may not contribute a significant impact.

**Figure 2.**
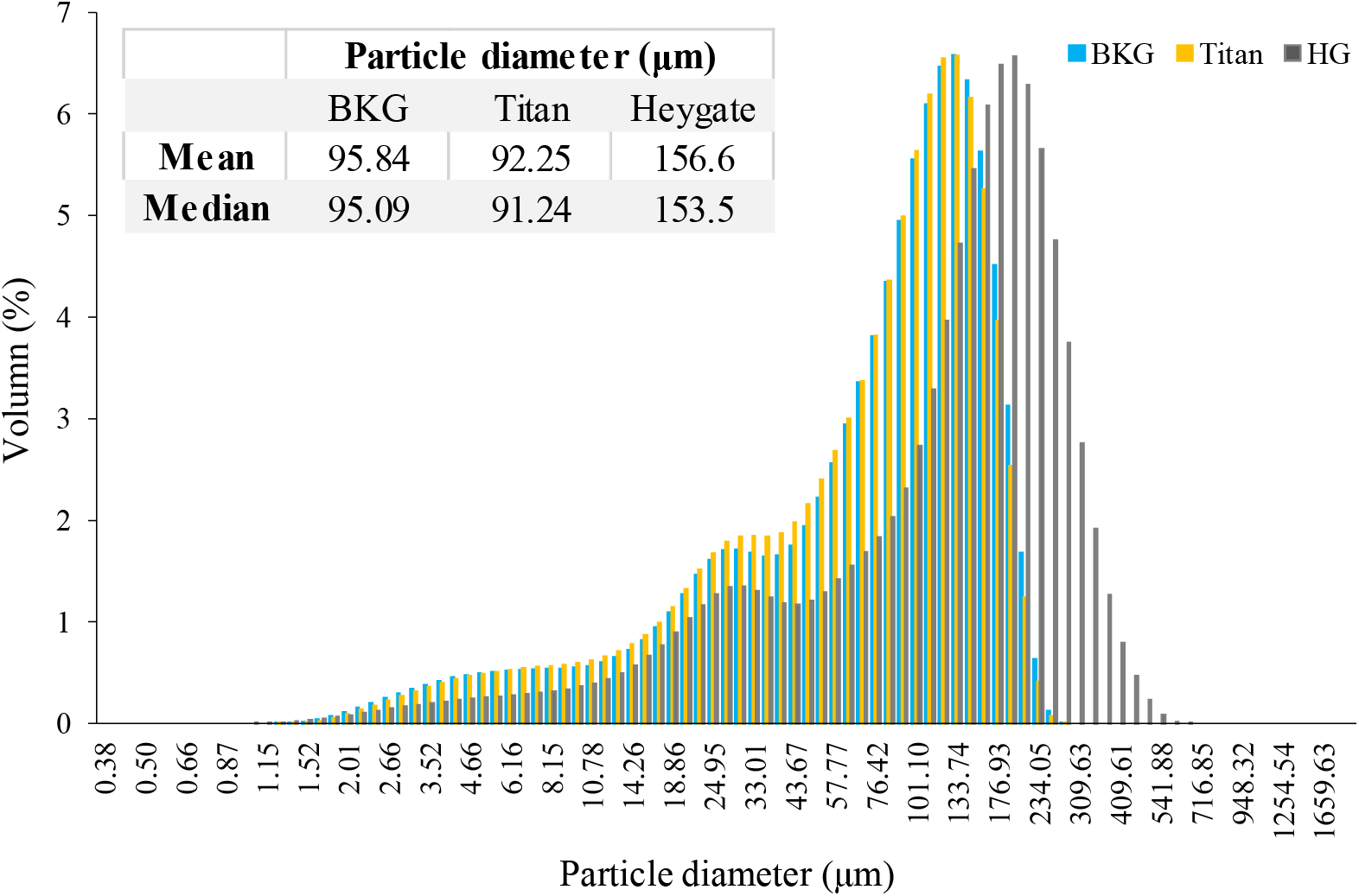
Particle size distribution for different wheat flours by using SLS. Blue bars, yellow bars and grey bars represent BKG, Titan and HG wheat flours respectively. The inset table shows the Mean and Median particle sizes for each wheat flour.

Wheat flours were analysed using XRD to investigate the level of starch crystallinity. As presented in Table 1, the relative crystallinity of BKG (39.9%) was slightly higher than Titan (38.2%) and HG (36.8%). The level of relative crystallinity is influenced by a number of factors: size of crystal, quantity of crystalline regions, double helices, orientation of samples, interaction between double helices, and even due to the interplay of these factors (Miao, Jiang, & Zhang, 2009; Song & Jane, 2000). Only relatively minor differences were observed in the relative crystallinity between the samples.

**Table 1.** Comparison of relative crystallinity and gelatinisation for 3 different wheat flours.

|  | Crystallinity (%) | Gelatinisation of starch |  |  |
| --- | --- | --- | --- | --- |
|  |  | Onset Temp (°C) | Peak Temp (°C) | Enthalpy (J/g) |
| **BKG** | 39.9 ± 1.07 | 53.65 ± 0.33 | 61.11 ± 0.10 | 4.43 ± 0.15 |
| **Titan** | 38.2 ± 0.43 | 53.05 ± 0.24 | 60.72 ± 0.08 | 4.94 ± 0.13 |
| **HG** | 36.8 ± 0.37 | 54.31 ± 0.03 | 60.59 ± 0.03 | 4.93 ± 0.08 |

The thermal behaviour and gelatinisation of the different wheat flours were investigated using DSC. The gelatinisation enthalpy and temperature is indicative of the relative amount and stability of the crystallites present in the starch (Warren, Gidley, & Flanagan, 2016). The results shown in Table 1 illustrate that the onset and peak temperatures and enthalpy of starch gelatinisation did not differ significantly between the wheat flours, except the gelatinisation enthalpy for BKG which was slightly lower (4.43 J/g) than other samples (4.49 J/g for Tian and 4.93 J/g for HG). Taken together, the results of the XRD and thermal analysis indicate that there were only minor differences in starch crystallinity between the different flours.

### 3.3 Aqueous wheat flours extracts are capable of reducing the digestion of purified wheat starch

To further investigate differences in starch digestion between the wheat flours, native starch was extracted from each flour and its digestibility was evaluated. As Figure 3 A shows, the digestibility of the extracted starches was not significantly different between the different wheat flours, as would be expected from their similar thermal and crystalline properties. This indicates that the differences in digestibility observed between the flours in Figure 1 were not due to intrinsic differences in starch digestibility. The starch digestibility of the control group and the gastric group was almost identical, and these results are similar to results for the control wheat starch shown in Figure 1. This is also consistent with the study of Zou et al. (2015) which showed that for Durum wheat there was no significant differences in starch digestibility between purified starch treated with and without pepsin. This provides strong evidence that the differences in starch digestibility between the wheat flours is as a result of the protein component in the flour. To test if the soluble albumin fraction or the insoluble gluten proteins were having the largest effect on starch digestion, the digestion of pure wheat starch was tested in the presence of aqueous extracts of the wheat flours.

**Figure 3.**
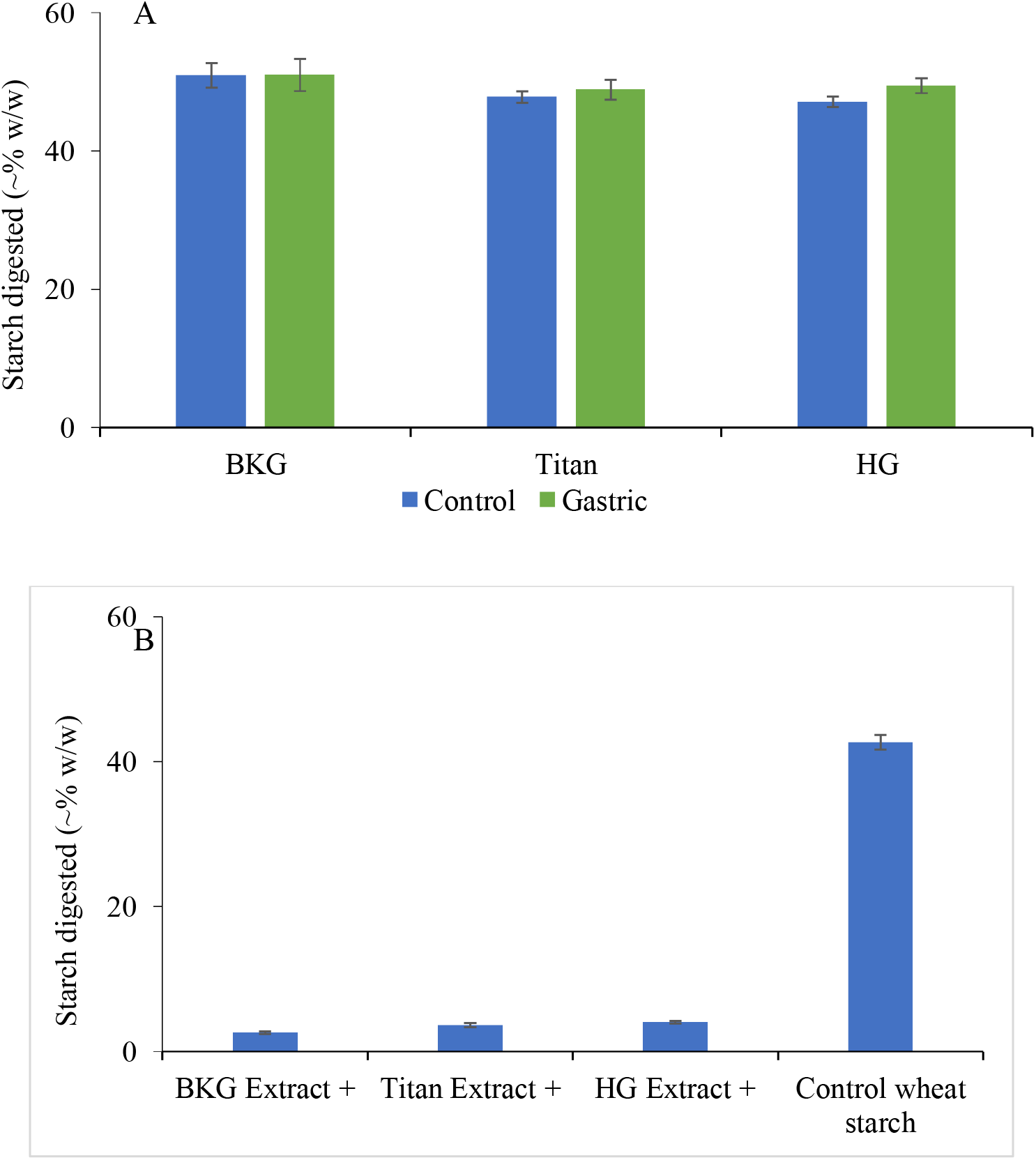
Results of starch digestion for starch extracted from wheat flours (Figure A) and the control wheat starch with the addition of wheat flour extracts (Figure B). In Figure A, BKG, Tian and HG represent the extracted starches corresponding to each of the wheat flours respectively. In Figure B, purified wheat starch is digested in the presence of aqueous extracts of each of the three wheat flours.

Results are shown as Figure 3 B. The starch digestibility of the control wheat starch was dramatically decreased with the addition of each of the aqueous extracts from the wheat flours. There was less than 5% starch digested with each of the extracts, whereas 42.7% of the starch was digested for the control wheat starch without addition of extracts. L. Li et al. (2022) reported inhibition of starch digestion caused by adding extracts from a range of cereals, such as sorghum, highland barley, barley and quinoa. Extracts from wheat were also used in the study of Franco et al. (2005) to supress activities of two different α-amylases. These studies had clearly pointed out that many aqueous cereal flour extracts would be able to inhibit starch digestion which has been hypothesised to be due to the presence of proteinaceous α-AIs. Therefore, we used proteomics methods to provide a more detailed characterisation of the proteins present in the aqueous extracts, and how the change as a result of gastric digestion.

### 3.4 Characterisation of glutenin’s, gliadins and proteinaceous α-AIs in the supernatants of wheat flours before and after gastric phase

Wheat flours were prepared separately as control and gastric digestion groups as described previously. The gastric group was treated with pepsin before collecting supernatant, whereas the control was not treated with pepsin. As Figure 4 shows, over 2000 proteins were detected by MS for each of 3 different wheat flours. Among all detected proteins, there were around 200 proteins significantly different in abundance between the control and gastric group. In Figure 4 the most relevant proteins have been indicated. A full list of proteins and NCBI codes is provided in Supplementary Table 1.

**Figure 4.**
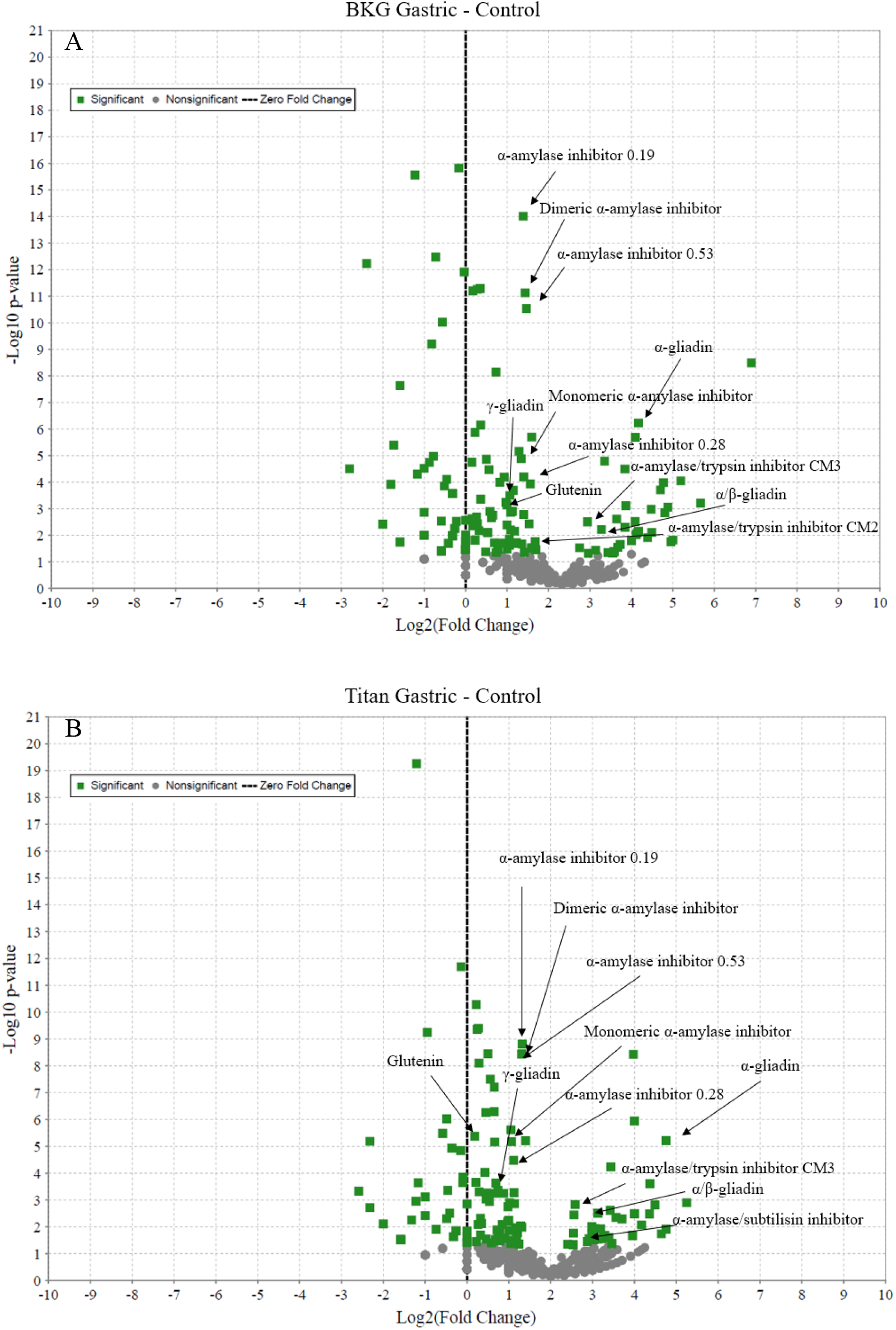
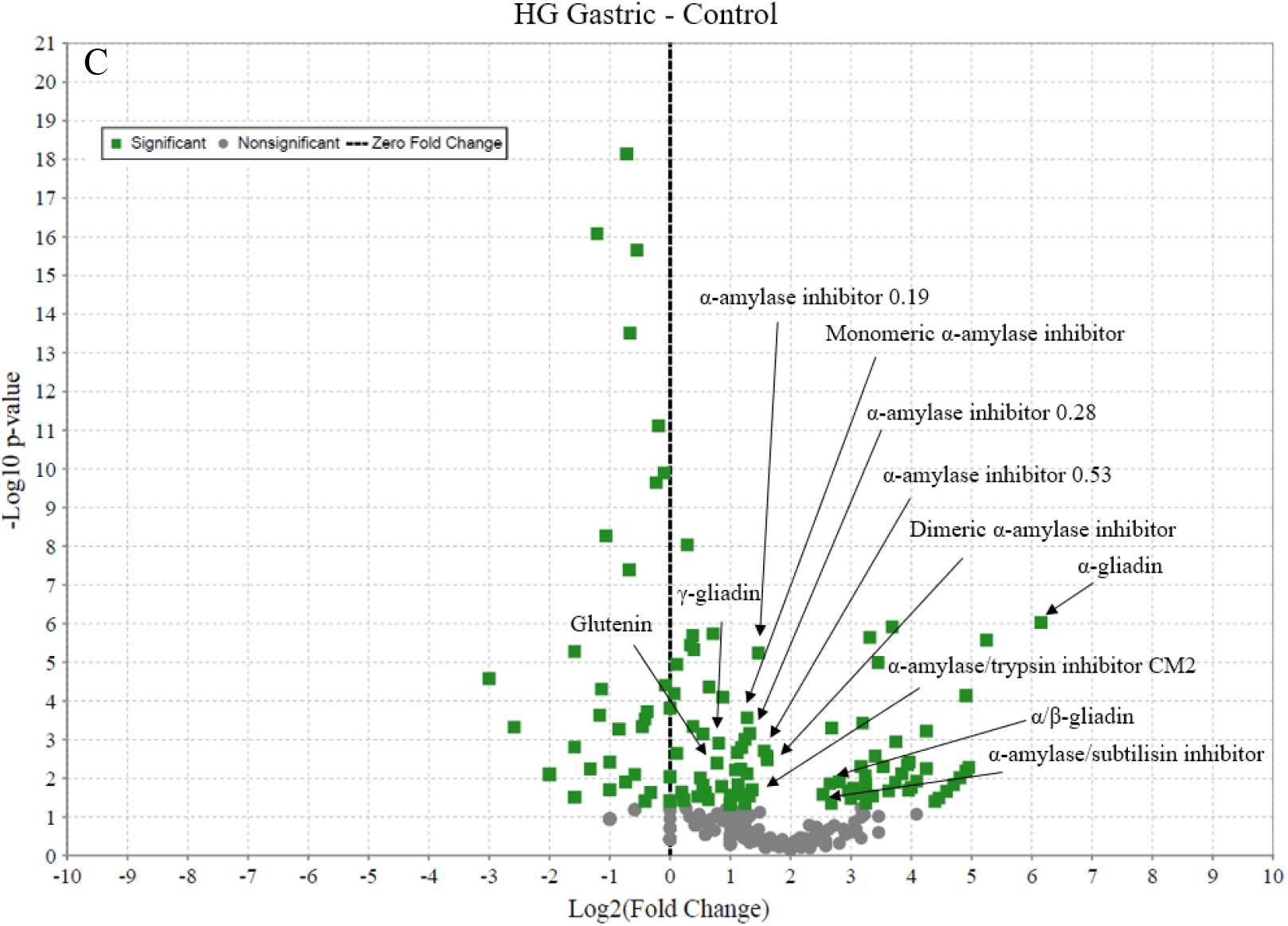
Volcano plots of proteins identified in supernatants of 3 different wheat flours treated (the gastric group) or not treated (the control group) with pepsin. Figure A, B, C are the results respectively for BKG, Tian and HG. The x axis is the measure of the strength of statistical signal, the data points with low p-values (high in significant) locating toward the top. The y axis is a measure of the statistical significance of the signal. Data points on the left side of the black dash line (zero-fold change) are proteins that higher in the gastric group, whereas data points on the right side are proteins higher in the control group.

Several proteinaceous α-AIs and gluten derived proteins were found significantly higher in the control group than in the gastric group. Such as α-amylase inhibitor 0.19, α-amylase inhibitor 0.53, α-amylase inhibitor 0.28, monomeric/dimeric α-amylase inhibitor, α-amylase/trypsin inhibitors, α-amylase/subtilisin inhibitor, gliadins and glutenin (component proteins of gluten). The soluble proteinaceous α-AIs identified were consistent with findings in the research of Salt, Robertson, Jenkins, Mulholland, and Mills (2005). All proteinaceous α-AIs found in wheat flour extracts were categorised as cereal type or CM type (inhibitors found in chloroform/methanol extracts of wheat) α-AIs, expect the α-amylase/subtilisin inhibitor which was classified as a Kunitz type (H. Li, Zhou, Zhang, Fu, Ying, & Liu, 2021; Svensson et al., 2004).

Proteinaceous α-amylase inhibitor 0.19, α-amylase inhibitor 0.53 and α-amylase inhibitor 0.28 were named according to their gel electrophoretic mobility of 0.19, 0.53 and 0.28 respectively to bromophenol blue (Feng, Richardson, Chen, Kramer, Morgan, & Reeck, 1996). The inhibitor 0.19 and 0.53 are acting as homodimers, whereas inhibitor 0.28 is a monomer. These inhibitors have very different preferences for α-amylases. The inhibitor 0.19 inhibits a broad range of α-amylases in mammals and insects and the inhibitor 0.28 is more active against α-amylases from insects than mammalian amylases, whereas the inhibitor 0.53 is specifically prefer α-amylases from insects (Svensson et al., 2004). The inhibition mode of these on porcine pancreatic α-amylase is non-competitive when starch is used as a substrate (Islamov & Fursov, 2007).

The α-amylase/trypsin inhibitors are actually a family of proteins (CM2, CM3 and CM16 were detected in this study), which inhibit α-amylases from both porcine pancreas and insects by binding to the active site of amylase, hindering contact between substrates and amylases and weakening the affinity of α-amylases to substrates (H. Li et al., 2021; Saxena, Iyer, & Ananthanarayan, 2010). The Kunitz type inhibitor - α-amylase/subtilisin inhibitor was widely found in cereals including barley, rice and wheat. Barley α-amylase/subtilisin inhibitor was the most studied which had been reported to specifically inhibits barley α-amylase 2 (H. Li et al., 2021; Svensson et al., 2004). The wheat α-amylase/subtilisin inhibitor has not been well studied, and it was reported to have 92% similarity to barley α-amylase/subtilisin inhibitor in amino acid sequences (Franco et al., 2004; Wisessing & Choowongkomon, 2012). This implies that wheat α-amylase/subtilisin inhibitor might also has preferences to inhibit wheat endogenous α-amylases. However, the Kunitz type inhibitors from legumes have been reported to be able to inhibit α-amylases from mammals and insects as well as plant α-amylases (Alves et al., 2009).

As shown in Figure 4, there were significant reductions in the abundances of all of these amylase inhibitors as a result of gastric proteolysis for all three of the wheat flours tested. Therefore, this data indicates that the reduction in abundance of proteinaceous α-AI inhibitors may explain larger differences observed between wheat flour digestion with and without a gastric phase, and as indicated in Figure 1, the incomplete proteolysis of these inhibitors (which were still identified by proteomics following the gastric phase) may impact wheat flour digestion and explain differences in digestibility between wheat flours from different sources with different protein contents.

In addition, there were also significant reductions in some glutenin and gliadin proteins. Wheat gluten had been extensively studied and indicated as not just a physical barrier to digestive enzymes but an effective inhibitor that binds to α-amylases (Bhattarai et al., 2016; Chen, He, Zhang, Sun, Liang, & Huang, 2019). However, in this aqueous extract, the glutenin and gliadin proteins were found at a much lower abundance than the α-AI’s, so may be playing a lesser role.

## 4. Conclusion

In this study, the role of wheat proteins and grain hardness in controlling starch digestibility was explored. Using wheat flours with different hardness and protein content, we demonstrated that wheat flour was significantly more slowly digested than purified starch, and that this effect was significantly reduced by gastric digestion and proteolysis. While grain hardness impacted on flour particle size, it did not have a large effect on starch digestibility. We then identified that an aqueous extract of wheat flour could dramatically slow starch digestion. Using proteomics, we identified that the aqueous extract contained a large number of proteinaceous α-AI’s which were only partially degraded by gastric protein digestion. Therefore, we conclude that soluble wheat proteins from both hard and soft wheats, particularly α-AI’s, can have a significant effect in modulating starch digestion in wheat foods, despite gastric protein digestion.

## Abbreviations

GI: glycaemic index
BKG: Baker’s green wheat flour
Titan: Titan wheat flour
HG: Heygates plain all-purpose wheat flour
α-AIs: α-amylase inhibitors
TTS: Total starch content
RTS: The rapid total starch method
PBS: Phosphate buffer
PAHBAH: p-hydroxybenzoic acid hydrazide
DSC: Differential Scanning Calorimetry (DSC)
XRD: X-ray diffraction
SLS: static light scattering
MS: Mass Spectrometry and Proteomics
SDC: sodium deoxycholate
SKCS: single-kernel characterization system
CAM: carbamido-methylation (CAM)

## Acknowledgements

We acknowledge the technical assistance of Bertrand Leze (University of East Anglia) with XRD analysis and Gerhard Saalbach (John Innes Centre) with proteomics analysis. We acknowledge Jan Delcour, Kurt Gebruers, Elien Lemmens, Leonardo Mulargia and Stijn Reyniers (KU Leuven) for helpful discussions in the preparation of this manuscript. FJW also gratefully acknowledges the financial support of the BBSRC Institute Strategic Programme Food Innovation and Health BB/R012512/1 and its constituent project BBS/E/F/000PR10345. This EIT Food activity has received funding from the European Institute of Innovation and Technology (EIT), a body of the European Union, under Horizon Europe, the EU Framework Programme for Research and Innovation. The report does not reflect the views of the European Union.

